# Hotspot prioritizations show sensitivity to data type

**DOI:** 10.1101/685735

**Authors:** Kari EA Norman, Ethan P White

## Abstract

Prioritizing regions for conservation is essential for effectively allocating limited conservation resources. One of the most common approaches to prioritization is identifying regions with the highest biodiversity, or hotspots, typically using global range map data. Range maps are readily available at large scales for an array of taxa, but are also known to differ from local-scale survey data in the same regions. We examined how prioritizations may differ between range map and survey data using the North American Breeding Bird survey (BBS) and BirdLife International range maps as a case study. Hotspot prioritizations were generated for species richness and the richness of rare species at two scales.

Total species richness patterns differed substantially between data types with at most a 41% overlap in identified hotspots. Some regions had few or no hotspots for one data type and a significant number for the other. Hotspots for rare species were more similar across the data types with 44% overlap at the larger scale. Future efforts to prioritize areas for conservation should consider differences between local-scale survey data and range maps, match data to the scale of interest, and develop methods to better downscale range map-based prioritizations to the scale of conservation decisions.

## INTRODUCTION

A key component of allocating limited conservation resources is identifying high priority regions for protection or management. Doing this effectively is essential to address threats to natural systems and the services they provide, given growing human populations, rising global temperatures, and widespread land use change. Approaches for systematically identifying the most important areas to conserve is broadly referred to as conservation prioritization^1^.

Conserving biodiversity is both a goal of conservation and a metric associated with other positive conservation outcomes^2^. As a result, many conservation prioritization analyses focus on maximizing the number of species (i.e., species richness) in a given area to be conserved. Other desirable conservation criteria include endemism (the number of species occurring only in a particular area), vulnerability (species designation as threatened or endangered), and level of threat (likelihood of future habitat loss).

Myers’ (2000) now-classic paper established biodiversity-based conservation prioritization with a global assessment that identified 25 global biodiversity hotspots based on vascular plant endemism and threat^3^. Building on this work, hotspot prioritizations have been created for a number of different taxa^4–7^, compared to current reserve networks to evaluate their effectiveness in protecting biodiversity^8–11^, and used to assess the scope of human impact on biodiversity centers^12^.

In addition, recent developments in software and methods (e.g., Zonation^13^ and Marxan^14^) provide improved prioritizations that incorporate greater levels of ecological and human complexity. Such advances have transformed conservation prioritization from a technique exclusively used in global-scale categorizations to a viable tool for local managers. Managers now use conservation prioritization to inform decisions such as the allocation of conservation money and expansion of local reserve networks^1^.

As use of conservation prioritization proliferates, a largely unacknowledged methodological divide has emerged between studies using two distinct kinds of data. Large-scale hotspot analyses rely almost exclusively on geographic range map data^3,4,7,8,15^. These data are heavily informed by expert opinion and potential habitat^16^. The relatively low cost of that information means range maps are accessible for a wide array of taxa at continental to global scales. However, range map data have two potential weaknesses: 1) typically, they are temporally static; and 2) they reflect biodiversity at spatial scales of nearly 2×2 degrees (∼40,000 km^2^)^16^. In contrast, managers working at smaller scales typically use survey data. Though costly to collect, these data provide direct observations of species richness or abundance in a particular region. These regions are often much smaller than the 2×2 degree grid cells approximated using range map data.

Despite this dichotomy between range map and survey-based approaches, there have been no analyses to examine how differences in the underlying data influence the regions prioritized for conservation. While range map suitability for use in IUCN classification has been questioned^17^, and comparisons of range map and survey data for biodiversity patterns more generally show significant disparities^18^, it is unclear how those discrepancies impact prioritization results.

Here we explore the sensitivity of biodiversity hotspots, the simplest conservation prioritization analysis, to the type of data (surveys or range maps) used in the analysis. We do this by combining data from a continental scale survey of breeding landbirds with data from the most widely used digital range maps. We compare the hotspots identified by these two data sources at two scales using both total species richness and the richness of rare species and discuss the implications of the results for future conservation prioritization efforts.

## METHODS

### Data Sources

We compared biodiversity patterns of North American breeding landbird species based on survey and range map data. Digital breeding range maps were obtained from Birdlife International^4,19^. Survey data were from 2,769 routes of the North American Breeding Bird Survey (BBS; acquired using the Data Retriever^20^), collected from 2005-2015^21^. Each route is 24.5 miles long and surveyed annually in June. Three-minute point counts are made along the route every 0.5 miles, in which every bird seen or heard within 400 m is recorded. Difficult to survey groups including waterbirds, shorebirds, nightjars, and owls were excluded from survey and range analyses.

Birds with less than 50% of their range in the survey area (based on range map data) were also excluded to prevent boundary effects for maps of rare species. Rarity was determined by the percentage of sites occupied for each species, with fewer occupied sites considered more rare. Number of occupied sites only acts as an appropriate surrogate for range size when the survey area covers the entire range. For example, species occurring primarily in Mexico with a portion of their range along the US-Mexico border would be consistently classified as rare despite having potentially large unsampled ranges. These species were excluded from both richness and rarity analyses to allow for comparison across analyses, but richness maps retaining excluded species have only marginal differences (Supplement). A complete list of species excluded from analysis can be found in the supplement.

### Analysis

The first step in comparing prioritizations based on survey and range map data was to ensure a fair comparison between inherently different data types and methodological approaches. Survey data are discrete point estimates of richness at each site. In contrast, range maps data are typically aggregated into cells and richness patterns analyzed across those cells. To accommodate both standard approaches we compared survey and range map based prioritizations at both site and cell levels for total species richness and the richness of rare species. Since all range maps (except those previously discussed) were included in the analyses regardless of whether the species was observed in survey data, total range richness is higher for range map data than survey data.

Point richness estimates for both data types were made at the starting position of each BBS survey route. For survey data, estimates were the number of species recorded at any time on each route over the ten-year survey period (2005-2015). All sites were included regardless of the number of times they were sampled to most closely reflect range maps, which are a representation of occurrence. The majority of sites (69%) were surveyed more than half of the years in the survey period. Range map estimates were calculated by counting the number of individual species ranges intersecting with each point. Cell level richness was calculated for 100 km^2^ cells across North America^22^. This is a typical cell area that accounts for range map resolution^16^ while still producing a large number of cells for analysis.

Maps for rare species were created using the same methods for site and cell level richness estimates. We classified a species as rare when it occurred at a proportion of sites less than the median proportion of site occurrence^4^, which was less than 4% or 5% of sites for survey and range data respectively. As previously discussed, the rarity proportion is sensitive to variation in the intensity of spatial sampling, as a species could be considered rare due to a lack of sampling locations within its range. Richness estimates for rare species were therefore based on a subset of sites adjusted to have consistent sampling intensity across the study area. This subset was made up of three randomly selected sites from within 100 km^2^ cells across North America. Cells containing less than three sites were excluded. This combination of cell size and number of samples was sufficient to address the bias while retaining a sufficient percentage of the data for a meaningful analysis.

The sites or cells with species richness in the top 5% were identified as hotspots, a commonly used cutoff for such analyses^4,16^. Hotspot locations based on survey data were compared to those based on range map data. The percent of hotspots that were shared between range map and survey based approaches was calculated by direct site-to-site and cell-to-cell comparisons.

All code for this project is available on GitHub (https://github.com/weecology/diversity-conservation) and archived on Zenodo (https://doi.org/10.5281/zenodo.3258263). Data for Breeding Bird Survey is downloaded by this code. Range map data can be obtained from http://datazone.birdlife.org/species/requestdis.

## RESULTS

### Richness

Maps of hotspot locations for species richness showed both similarities and notable differences at the site and cell level (Figure 1). At the site level, survey data show hotspots across the northeastern United States to the Great Lakes region up into Lake Winnipeg, and stretching down to southern states. Smaller hotspot contingents are also scattered across the Rockies, the Colorado Plateau, and California. Range map data show generally less dispersed priority regions likely due at least in part to the much higher spatial autocorrelation inherent in range map data^18^. Heavily concentrated priority areas occur primarily in the northern Rockies, the Colorado Plateau, and the Great Lakes Region into Lake Winnipeg. Smaller hotspots were also identified in the mountains of northern California and New York. In general range map data show a much higher concentration of hotspots in the Rocky Mountains and a general absence of the hotspots identified by survey data in the Appalachian Mountains. As a result, range map analyses gave higher priority to regions in western North America, while analyses of survey data gave higher priority to eastern North America. Hotspot similarity was low at only 22% (Figure 3).

**Figure 1.**
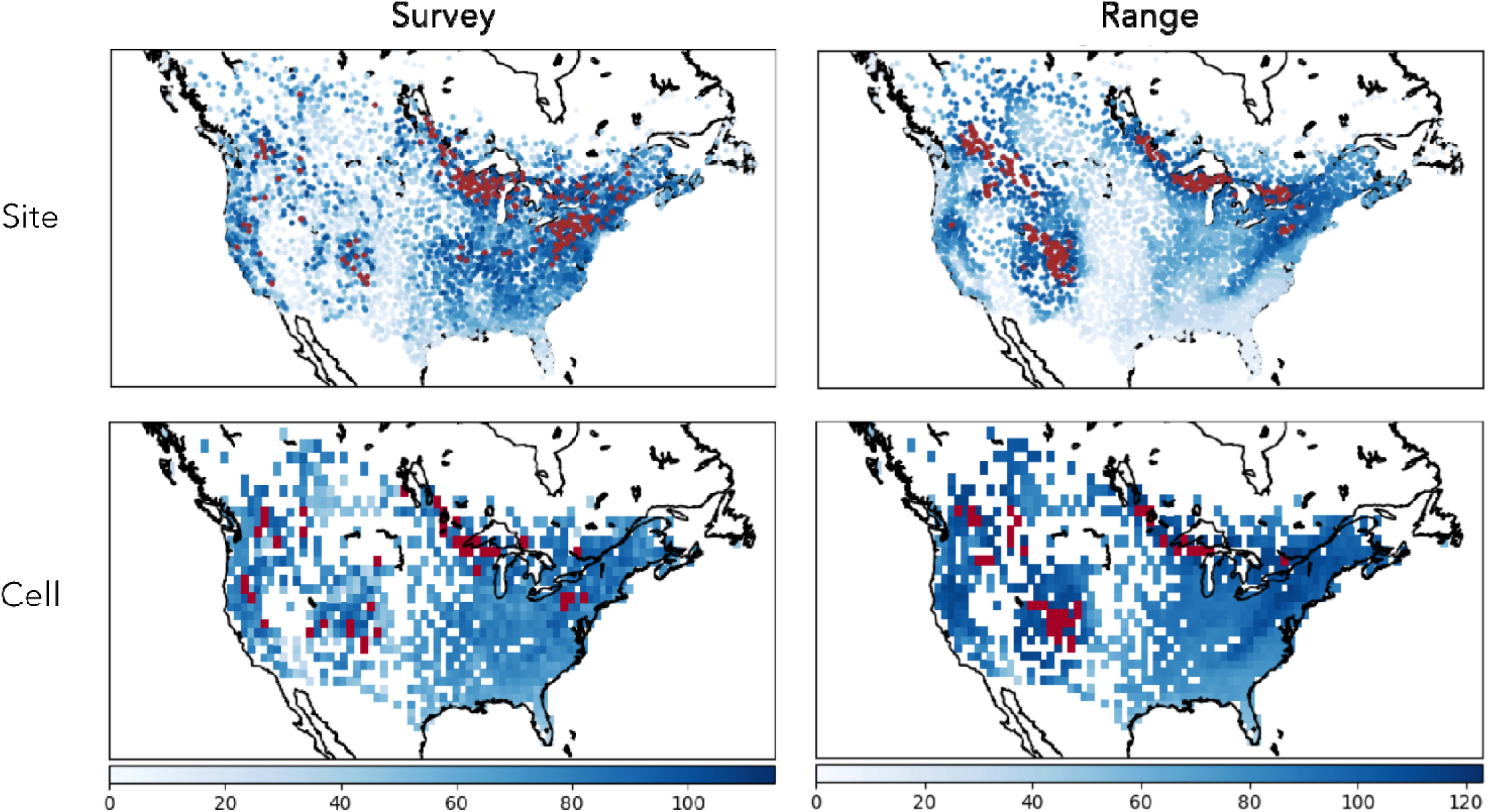
Maps of species richness based on survey and range map data at the site and cell level. Cell level richness is a count of species recorded at any site within the cell. Darker regions correspond to higher biodiversity, with hotspots (richest 5%) marked in red.

At the cell level, hotspots for survey data remain largely concentrated in the Great Lakes region and New England. Scattered hotspots in the Rockies, California and the Colorado Plateau aggregated to maintain hotspot representation in those areas while hotspots in the Midwestern and southern states disappeared. Range map data show similar hotspot distributions to those at the site level, though the small number of hotspots in the central Great Lakes region and northern Californian disappeared. Concentrations of hotspots remained around Lake Winnipeg, Lake Superior, Lake Ontario, the northern Rockies and the Colorado Plateau (Figure 1). This resulted in more similar maps and greater hotspot similarity, 41%, between the survey and range map relative to the site level (Figure 3). However, range maps gave higher priority to the Colorado Plateau than the survey maps which prioritized the Great Lakes region and northern states more heavily.

### Rarity

Richness patterns of rare species show more congruence across level and data type (Figure 2, Figure 3). All four prioritizations identify hotspots along the western coast and the Colorado Plateau, with a larger concentration of hotspots on the Colorado Plateau for range maps and British Columbia for survey maps. Hotspots for rare species had 42% overlap at the site level and 44% at the cell level (Figure 3).

**Figure 2.**
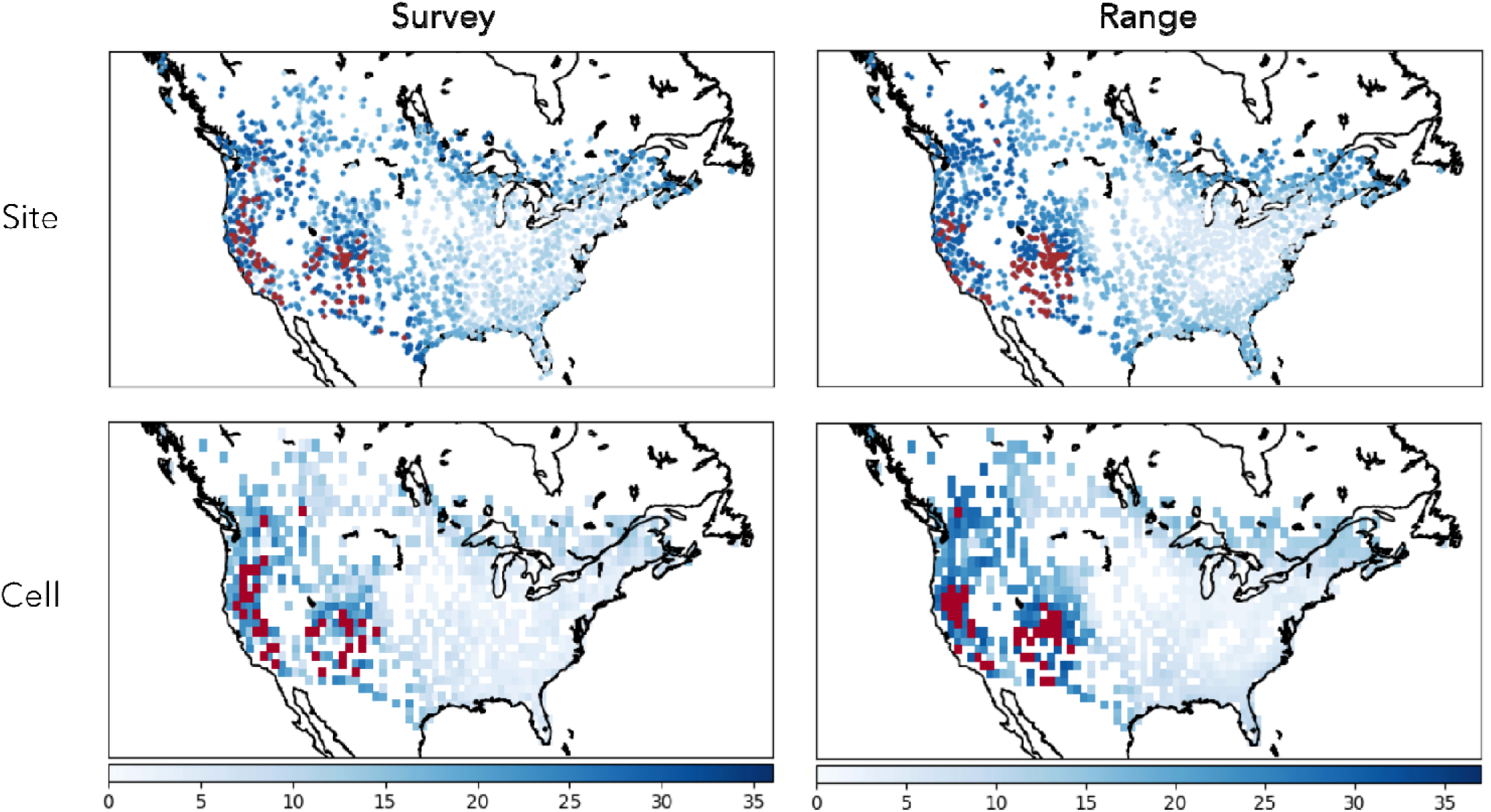
Maps of the richness of rare species based on survey and range map data at the site and cell level. Cell level richness is a count of rare species recorded at any site within the cell. Darker regions correspond with more rare species, with hotspots (richest 5%) marked in red.

**Figure 3.**
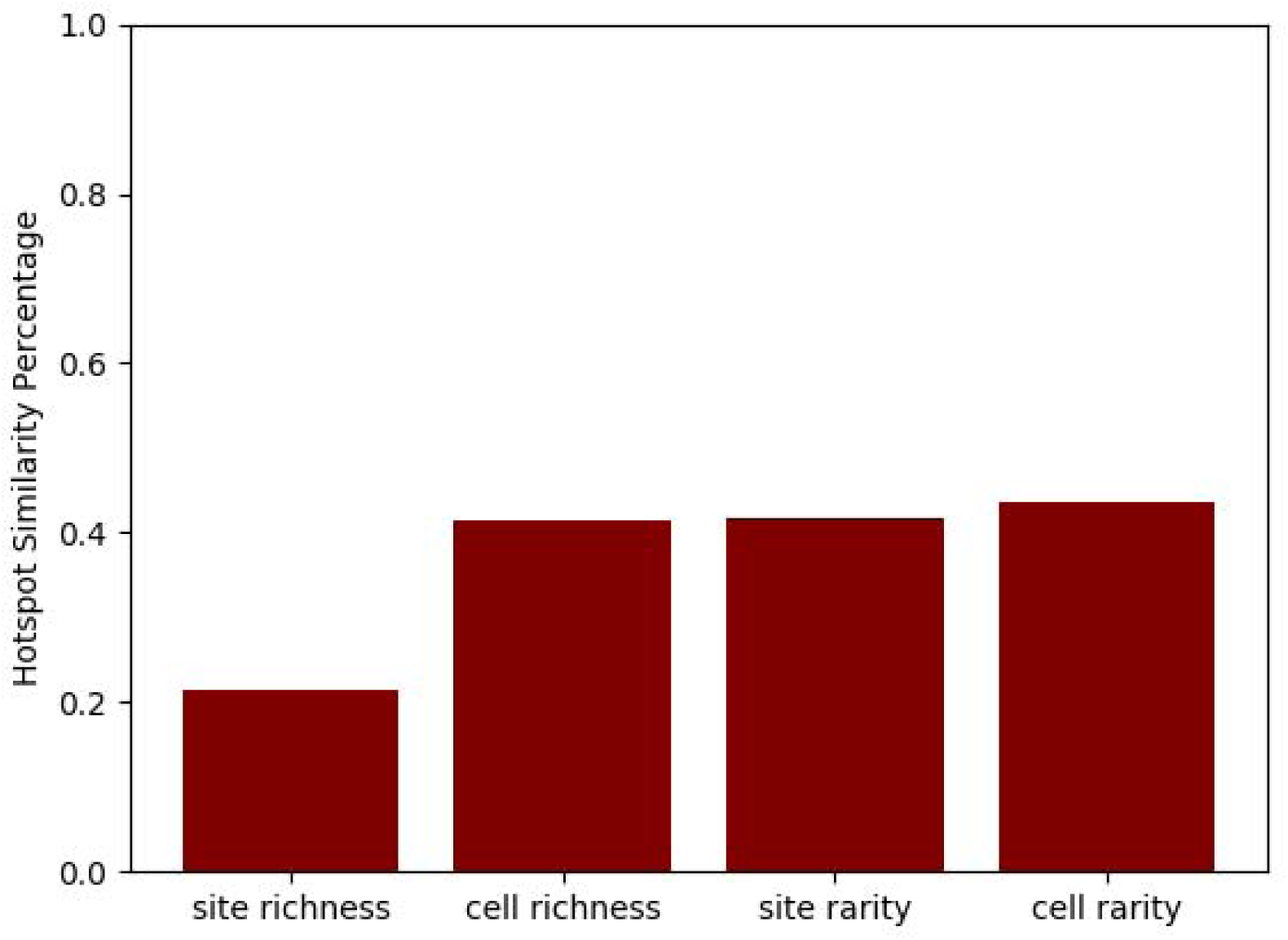
Percent hotspot similarity between range map and survey data types for richness and rare species at the site and cell level.

## DISCUSSION

Comparisons of conservation priority areas based on survey and range map data show both similarities and notable differences between data types. Some of the discrepancies are substantial, particularly when evaluating total species richness at the site level. While some hotspots were in qualitatively similar regions for both data types, direct site comparison showed only a 22% overlap (Figure 3) and there were large regions for which there was high biodiversity for one data type and low biodiversity for the other. Aggregation to the cell level increased regional consistency qualitatively but still resulted in only 42% overlap between identified hotspots for the two data types (Figure 3). Even for regions prioritized by both data types, maps differ notably in which regions are most heavily emphasized. The Colorado Plateau has the largest concentration of hotspots for range maps and only a few for survey maps, while the reverse is true of the northeast with many hotspots for survey data and few hotspots for range map data. These discrepancies highlight the necessity for understanding the drivers of difference between data types and identifying the appropriate contexts for each to ensure the most informed conservation decisions. This need is especially pressing given uncertainty in how discrepancies may propagate through the increasingly complex prioritization analyses that are becoming more common.

Priority areas based on rare species richness were more congruent (Figure 2) with 42% and 44% overlap for site and cell level comparisons respectively (Figure 3). Hotspot regions were qualitatively consistent across data type and level, with hotspots along the western coast and on the Colorado Plateau. Greater overlap in hotspots for rare species compared to overall richness could be attributed to the different drivers that total richness and richness of rare species respond to. Species richness patterns are driven primarily by species with wide ranges and therefore drivers such as area, habitat heterogeneity and productivity^23^. In contrast, the most important driver for rare species is topographic heterogeneity^23^. Topographic heterogeneity is also generally positively associated with range map richness but negatively associated with survey richness^18^. Since range map richness and the richness of rare species are both positively influenced by topographic heterogeneity, similarity in drivers offer one possible explanation for higher congruence between data types for rare species than for common ones. To use both data types effectively further work is needed to understand the drivers of consistencies and inconsistencies between data types.

In choosing a data type for future prioritizations, one approach is to match the scale of the data type to the scale at which decisions are being made. Our cell level analyses were performed at a resolution of approximately 1 degree (100 km^2^ cells), a resolution typical for range map-based prioritizations. However, this is a coarser grain than would typically be used for local management decisions. As described in Jenkins et al.^4^, an area of that size in some parts of the world contains multiple mountain ranges and valleys. Attempts to simply analyze range map data at a scale more appropriate for conservation (e.g., Jenkins et al.^4^) may be misleading, as the inherent resolution of most range maps by some assessments is only 2 by 2 degrees^16^. Analysis of range map data below this scale results in overestimates of biodiversity and distorted spatial patterns^16^.

The mismatch between the resolution of range map data and the scale of conservation questions leaves survey data as a natural replacement. The local scale of survey data captures biodiversity patterns more representative of reality at the scales important for conservation. However, the use of survey data does present challenges. Important considerations for survey data use include the difficulty in accounting for incomplete surveys due to imperfect detectability^24^ and the availability of high quality data for a variety of taxa and locations^25^. Methods developed to adjust for imperfect detectability show estimated richness patterns that are highly correlated with observed richness in birds, indicating that detectability likely does not have a large influence on spatial patterns of richness^26^. The increased availability of high-quality survey data through community science and large-scale government efforts means that using survey data in place of range map data is increasingly possible for a greater variety of taxa. Efforts like the National Ecological Observatory Network and burgeoning community science programs will assure that more data for survey based assessments is available in the future. The potential for survey data to improve the accuracy of biodiversity-based conservation indicates that its further availability is a worthwhile investment for the conservation community.

Despite the many benefits of survey data, conservation decisions cannot wait for more comprehensive data availability. It is therefore important to explore approaches for using currently available survey and range data as effectively as possible. Developments in single species and joint species distribution modeling offer promising approaches for filling geographic and temporal gaps in survey data using environmental information^27,28^. Spatial interpolation has been shown to perform well for smoothing spatial gaps in survey data^29^. Potential improvements in range maps include methods for downscaling range map data to more useful scales^30,31^, updating old maps to reflect changes such as range shifts and land use changes, and new approaches for addressing range map porosity^18^. Establishing relationships between richness drivers and congruence between data types, as previously described, will also play an important role in making methods for accurate range map data use possible. Development of new methods that integrate survey and range map data to inform occurrence extent would greatly improve use of all available species data.

Our findings underline the importance of understanding the implications of data type when prioritizing areas for conservation. Discrepancies in hotspot location and overall biodiversity patterns between data types give evidence of the current tradeoff between accuracy and availability in data type selection. Further exploration of biodiversity analyses’ sensitivity to data type is essential for partitioning the appropriate roles of each data type and ensuring the most effective conservation planning into the future.

## Acknowledgements

We thank the volunteers and researchers at USGS and CWS for collecting and making available BBS data, and likewise the researchers at Birdlife International. This research was supported by United States Department of Energy through the Computational Sciences Graduate Fellowship (DOE CSGF) under grant number DE-FG02-97ER25308 awarded to K.E.A.N, the Gordon and Betty Moore Foundation’s Data-Driven Discovery Initiative through grant GBMF4563 to E.P.W., and by the National Science Foundation through grant 1354563 to Allen H. Hurlbert and E.P.W.

