## Supplement for "Hotspot prioritizations show sensitivity to data type"

**Supplementary Materials**

*Species Exclusions*

Species that are poorly represented by BBS survey data were excluded from all analyses. For rarity analyses, rarity was defined by the proportion of sites in which a species was found. For species whose ranges extend beyond the survey boundaries, rarity would therefore be artificially inflated. Species for whom the ranges were not contained mostly (>50%) within the survey extent were therefore excluded from rarity analysis. For ease in comparison across figures they were also excluded from richness analyses. Figure 1 demonstrates the insensitivity of richness maps and hotspot locations to the exclusion of non-North American majority species, with almost near-identical qualitative patterns between maps with and without excluded species, and 90% and 83% overlap in hotspots comparing across exclusion for survey data and range map data respectively.

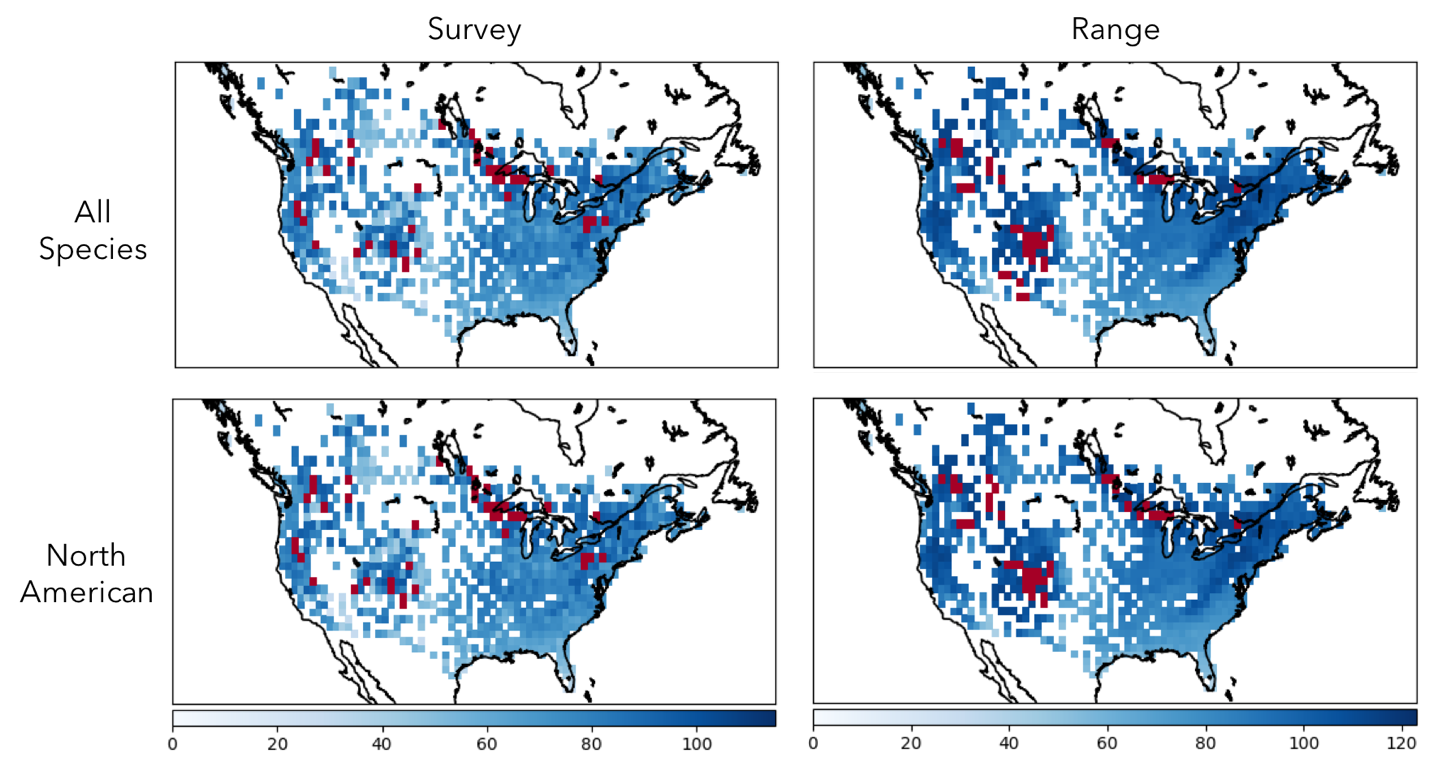

**Figure 1.** Comparison of richness maps with and without species whose ranges fall mostly (>50% of area) outside of North America.

Excluded Families:

Anatidae, Alcidae, Stercorariidae, Laridae, Sternidae, Rynchopidae, Scolopacidae, Recurvirostridae, Charadriidae, Haematopodidae, Jacanidae, Gaviidae, Gruidae, Aramidae, Rallidae, Anhingidae, Phalacrocoracidae, Pelecanidae, Fregatidae, Phoenicopteridae, Podicipedidae, Diomedeidae, Procellariidae, Hydrobatidae, Phaethontidae, Sulidae, Procellariidae, Tytonidae, Strigidae, Alcedinidae, Upupidae, Todidae, Momotidae, Coraciidae, Cinclidae, Threskiornithidae, Ciconiidae, Ardeidae

Additional Excluded Species:

Cypseloides niger, Chaetura pelagica, Chaetura vauxi, Aeronautes saxatalis, Eugenes fulgens, Lampornis clemenciae, Archilochus colubris, Archilochus alexandri, Calypte costae, Calypte anna, Selasphorus platycercus, Selasphorus rufus, Selasphorus sasin, Stellula calliope, Calothorax lucifer, Amazilia yucatanensis, Amazilia violiceps, Cynanthus latirostris, Coragyps atratus, Columba livia, Columba leucocephala, Streptopelia chinensis, Leptotila verreauxi, Columbina passerine, Scardafella inca, Streptopelia decaocto, Crotophaga sulcirostris, Geococcyx californianus, Coccyzus minor, Coccyzus americanus, Coccyzus erythropthalmus, Elanoides forficatus, Chondrohierax uncinatus, Elanus leucurus, Rostrhamus sociabilis, Parabuteo unicinctus, Buteo albonotatus, Buteo albicaudatus, Buteogallus anthracinus, Falco rusticolus, Falco femoralis, Caracara cheriway, Pandion haliaetus, Callipepla squamata, Cyrtonyx montezumae, Lagopus lagopus, Ortalis vetula, Pica hudsonia, Tyrannus crassirostris, Tyrannus melancholicus, Pitangus sulphuratus, Myiodynastes luteiventris, Myiarchus tuberculifer, Pyrocephalus rubinus, Camptostoma imberbe, Aphelocoma ultramarine, Cyanocorax yncas, Icterus graduacauda, Loxia curvirostra, Loxia leucoptera, Carduelis hornemanni, Carduelis flammea, Calcarius lapponicus, Junco phaeonotus, Amphispiza belli, Aimophila botterii, Aimophila carpalis, Arremonops rufivirgatus, Pipilo fuscus, Cardinalis sinuatus, Riparia riparia, Vireo huttoni, Parula pitiayumi, Peucedramus taeniatus, Dendroica coronate, Myioborus pictus, Passer domesticus, Passer montanus, Cardellina rubrifrons, Motacilla flava, Anthus rubescens, Anthus cervinus, Toxostoma longirostre, Toxostoma curvirostre, Toxostoma crissale, Campylorhynchus brunneicapillus, Baeolophus wollweberi, Poecile sclateri, Auriparus flaviceps, Phylloscopus borealis, Polioptila melanura, Polioptila californica, Luscinia svecica, Picoides scalaris, Picoides arizonae, Picoides tridactylus, Melanerpes formicivorus, Melanerpes aurifrons, Melanerpes uropygialis, Colaptes auratus, Colaptes chrysoides, Melopsittacus undulates, Myiopsitta monachus, Brotogeris chiriri, Trogon elegans, Geranoaetus albicaudatus, Patagioenas leucocephala, Columbina inca, Phasianus colchicus, Pachyramphus aglaiae, Corvus imparatus, Aphelocoma wollweberi, Cyanocorax morio, Acridotheres tristis, Cyanecula svecica, Motacilla tschutschensis, Acanthis flammea, Amphispiza quinquestriata, Peucaea carpalis, Peucaea botterii, Melozone fusca, Passerculus guttatus, Icterus gularis, Leiothlypis crissalis, Setophaga pitiayumi, Sporophila morelleti, Dryobates scalaris, Leuconotopicus arizonae
